# Mutational analysis unveils the temporal and spatial distribution of G614 genotype of SARS-CoV-2in different Indian states and its association with case fatality rate of COVID-19

**DOI:** 10.1101/2020.07.27.222562

**Authors:** Ballamoole Krishna Kumar, Bakilapadavu Venkatraja, Kattapuni Suresh Prithvisagar, Praveen Rai, Anusha Rohit, Madhura Nagesh Hegde, Indrani Karunasagar, Iddya Karunasagar

## Abstract

Pan genomic analysis of the global SARS-CoV-2 isolates has resulted in the identification of several regions of increased genetic variation but there is absence of research on its association with the clinical outcome. The present study fills the vacuum and does mutational analysis of genomic sequence of Indian SARS-CoV-2 isolates. Results reveal the existence of non-synonymous G614 spike protein mutation in 61.45% of the total study genome along with three other mutations. Further, temporal variation in the frequencies of G614 genotype in the country is observed. The examination of the probable association of G614 genotype with COVID-19 severity shows that CFR G614 genotype in India is positively and strongly correlated. It appears that the clinical outcome of the COVID-19 cases in India are significantly and adversely affected by the increasing trend in the G614 genotype; which needs to be addressed combining both laboratory experiments and epidemiological investigations.

## Introduction

Much like the world, India is fighting against an exponential growth of COVID-19 that started with the first imported case reported in late January 2020 from the Indian citizen travelled from Chinese city of Wuhan (Potdar et al., 2020). As of 21^st^ July 2020, the country has recorded over 11, 55,190 positive cases involving 28,084 deaths due to COVID-19 (https://www.mygov.in/covid-19). The interim analysis of the genomic data on Indian isolates of SARS-CoV-2 shows the introduction of this virus to India from multiple sources such as China, Europe, USA, Canada and the Middle East (Potdar et al., 2020; Singh and Sharma 2020; Somasundaram et al., 2020). SARS-CoV-2 has a genome size of 29.9 kb which encodes four major structural proteins [spike (S), envelope (E), membrane (M), as well as nucleocapsid (N) protein], 16 non-structural proteins [nsp1–16] and five to eight accessory proteins (Wu et al., 2020). Based on the currently available pieces of evidence, it is noteworthy that the severity of COVID-19 differs significantly depending on the geographical locations and this could be due to amalgamation of different factors including genetics of the virus.

All RNA viruses are prone to acquire an extremely high rate of mutation in the genomic region, which provides a selective advantage to the virus for transmission and virulence. However, coronaviruses have been reported to have genetic proof-reading mechanisms and therefore, nucleotide sequence diversity in SARS CoV2 has been found to be very low (Fauver et al., 2020). Nevertheless, pan genomic analysis of the global SARS-CoV-2 isolates has resulted in the identification of several genomic regions of increased genetic variation, which suggests that the virus underwent evolutionary changes and diversified during the process of geographical dissemination. Acquisition of non-synonymous D614G (Aspartate to Glycine) mutation in C-terminal region of the spike protein is one such evolutionary change observed in the SARS-CoV-2 genome and has become the most dominant genotype (>70%) by replacing D614G wild type strain in many countries and this mutation has been suggested to confer fitness advantage (Korber et al., 2020; 1. Biswas and Majumder 2020). In this study genomic sequence of Indian SARS-CoV-2isolates were examined for the temporal dynamics in the acquisition of D614G mutation and its probable relation to the case fatality rate among COVID-19 cases in India.

## Materials and Methods

A total of 1375 SARS-CoV-2genome sequences available from the different Indian states were retrieved from GISAID (https://www.gisaid.org/) database on 6^th^ July 2020. While retrieving the sequences the low coverage sequences with >5% Ns were excluded. It has been ascertained that all the retrieved sequences were mapped to reference genome of SARS-CoV-2deposited from Wuhan, China (GenBank Accession No. NC_045512.2) using Geneious Pro (V.8.0.5). The statistical data of daily reported cases and daily death toll (till June 30, 2020), case fatality rate (CFR) as per 7^th^ July 2020 were obtained from COVID-19 India website (https://www.api.covid19india.org/). The mutation analysis was performed using COVID-19 genome annotator as per the method described by Mercatelli et al., (2020). Mutations observed in >1.5% of the total sequences were tabulated. As this study is focused on the temporal dynamics in the acquisition of D614G mutation in the Indian SARS-CoV-2 isolates, its existence was further confirmed by manually analyzing the alignment created using Geneious (V.8.0.5). The D614G mutation was analyzed state-wise and in time series at 15 days interval from 1^st^ March 2020 to 30^th^ June 2020. The percentage of D614G mutation was then correlated with CFR. Also, the percentage of G614 mutant genotype and CFR were subjected to linear regression analysis using least squares method.

## Results

The study identifies 47 key mutations spread over different coordinates of Indian SARS-CoV-2isolates, which includes 25 non synonymous, 19 synonymous and 3 mutations were extragenic in nature (Table S1). The most frequently occurring mutations among the genomes studied are represented in Fig 1 with NSP3:F106F followed by S: D614G from spike protein, 5ÚTR:241, NSP12b: P314L subunit of RdRp and so on. Among these, acquisition of non-synonymous D614G (Aspartate to Glycine) mutation positioned between the receptor binding domain and S1/S2 polybasic protease cleavage site of spike glycoprotein were observed in the 61.45 % (n= 845/1375) of the total study genome. We have also reported the simultaneous occurrence of three other mutations such as 5’UTR: 241 C > T (61.01%, 840/1375), NSP12b: P314L (58.18%, 800/1375) and NSP3 F106F (3037 C > T) (62.03%, 853/1375) silent mutation among the SASRS-CoV-2isolates which showed D614G variation in the study. From here onwards, SARS-CoV-2isolates harboring these four distinctive mutations will be referred as G614 genotype.

**Fig 1.**
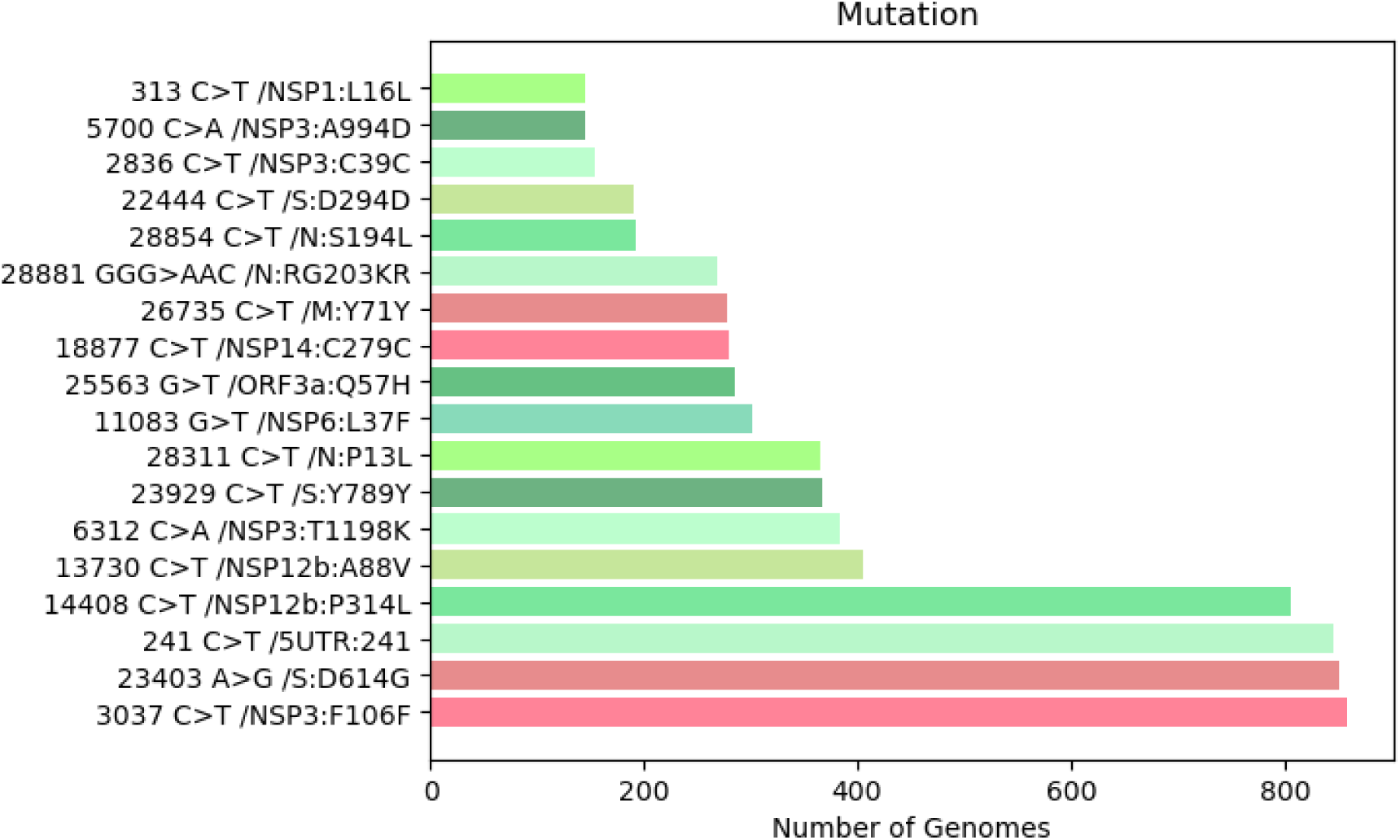
Analysis shows the most frequently found nucleotide variations (both synonymous and non-synonymous) in the Indian SARS-COV-2isolates.

The genomic leader sequence at 5’UTR plays an important role in the viral replication and gene expression during discontinuous sub-genomic replication of RNA viruses. Previous *in silico* analysis has shown that mutation of 5’UTR: 241 from C to T enhances the binding of Transactive Response DNA binding protein (TARDBP) to 5’-UTR of SARS-CoV-2 genome that could enable the virus to multiply within the host. (Mukherjee and Goswami, 2020). NSP12b: P314L is another prominent mutation observed among the Indian SARS-CoV-2isolates; found in very close proximity to the drug binding region in the hydrophobic cleft of RNA dependent RNA polymerase (Gao et al., 2020; Pachetti et al., 2020). RdRp is a key enzyme involved in the replication of SARS-CoV-2and exists as a complex of nonstructural protein 12 and 8 (NSP12 and NSP8); which is also a potential target for nucleotide analog antiviral inhibitors (Gao et al., 2020). 3037C>T is a silent mutation observed in the phosphoesterase, papain-like proteinase of SARS-CoV-2genome. Results obtained in this study are consistent with the previous findings where a simultaneous occurrence mutation in the different regions of the newly emerged mutant genotype of SARS-CoV-2has been reported (Korber et al., 2020). It is believed that, mutations in these structural and nonstructural proteins would have considerable impact on the replication of virus and their subsequent binding to the ACE2 receptor on the host cells. A spurt in the incidence of newly emerged G614 genotype was evident over wide geographical areas; afflicting an exceptionally high proportion of population. Studies have observed that there was dramatic increase in the incidence of SARS-CoV-2 carrying these distinctive mutations which may be due to the natural selection process (Korber et al., 2020; Pachetti et al., 2020). Based on data on RT-qPCR cycle threshold (CT) values, Korber et al., (2020) noted that G614 genotype infection leads to higher viral loads compared to D614, but no correlation with disease severity could be observed. On the other hand the functional study of G164 mutant utilizing retroviruses pseudotyped with G614 variant of spike protein showed greater infectivity potential compared to wildtype (Zhang et al., 2020).

Remarkably, none of the initial SARS-CoV-2 sequences from India had this distinctive D614G mutation; however, the first case of G614 genotype was detected in the early week of March from travelers arriving from European countries. From the analysis it is observed that there was a significant temporal variation in the frequencies of G614 genotype in the country as it contributed to 29.35% of the 109 genomes in March, 48.81% of the 338 genomes in April, 58.99% of the 656 available genome in May and 96.66% of 270 genomes in the month of June (Fig. 3). Our data indicates that in India, the transition from D614 to G614 genotype has grown progressively to become a dominant genotype and successfully established its niche in the country. Further, it is also evident that G614 genotype has a varying spatial distribution in the different Indian states; the highest percentage (93.04 %) being reported from the state of Gujrat followed West Bengal, Maharashtra, Madhya Pradesh, Orissa and Karnataka (Fig. 3). It is also noticed that, G614 genotype was not reported in some of the Indian states as represented in the Fig. 3 does not mean that it’s not prevailing because genome data from those regions are not sufficient to conclude its existence.

**Fig 2.**
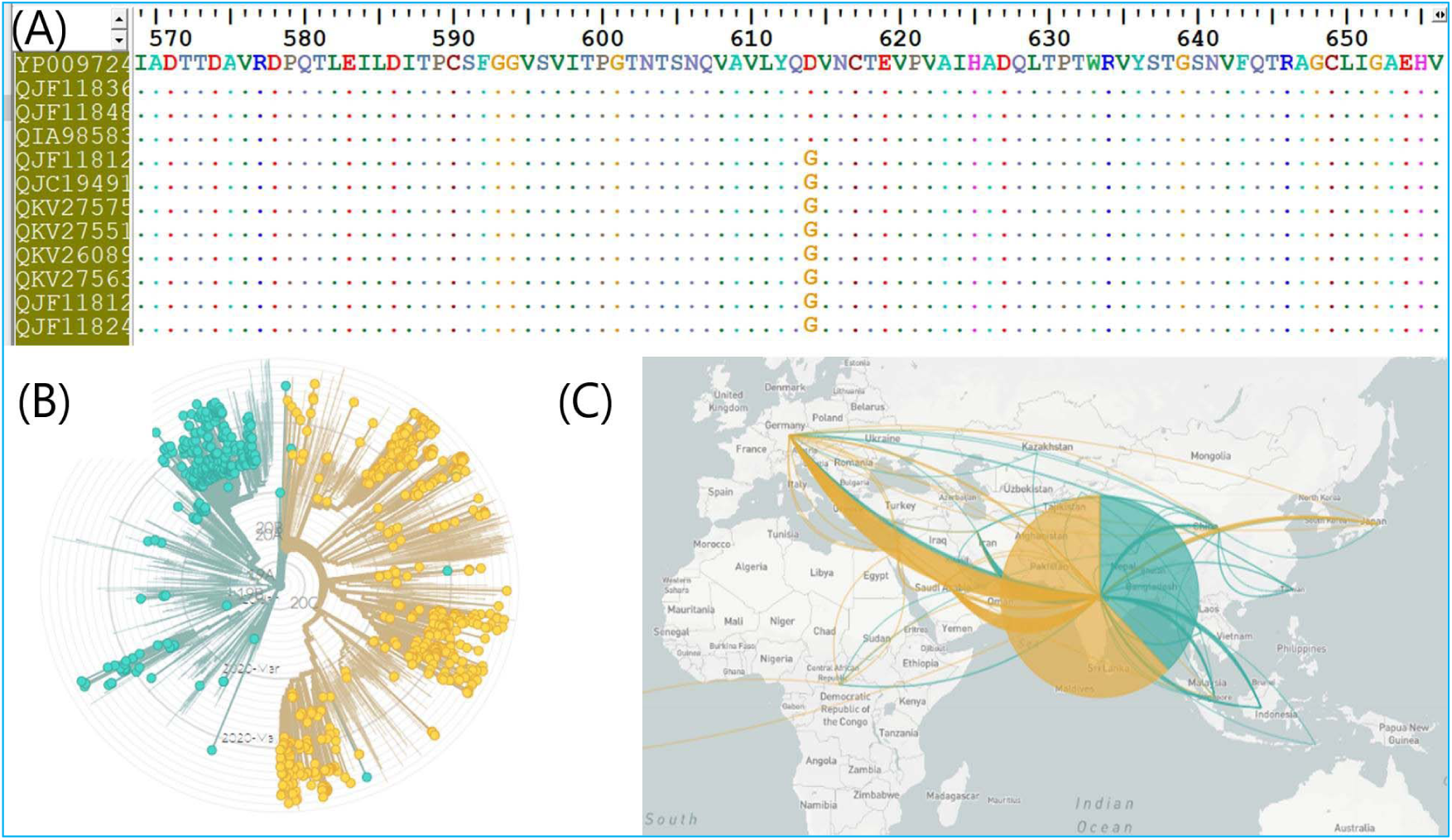
Analysis unveils presence of G614 genotype of SARS-CoV-2in Indian states. .Fig (A) Amino acid alignment of the S protein showing the D614G mutation sites among the selected SARS-COV-2genomes and (Fig B) separation of two major clades in the SARS-COV2-circulating in India as observed by phylogenetic analysis. (Fig. C) Transmission map showing the introduction of G614 genotype to India.

**Fig 3.**
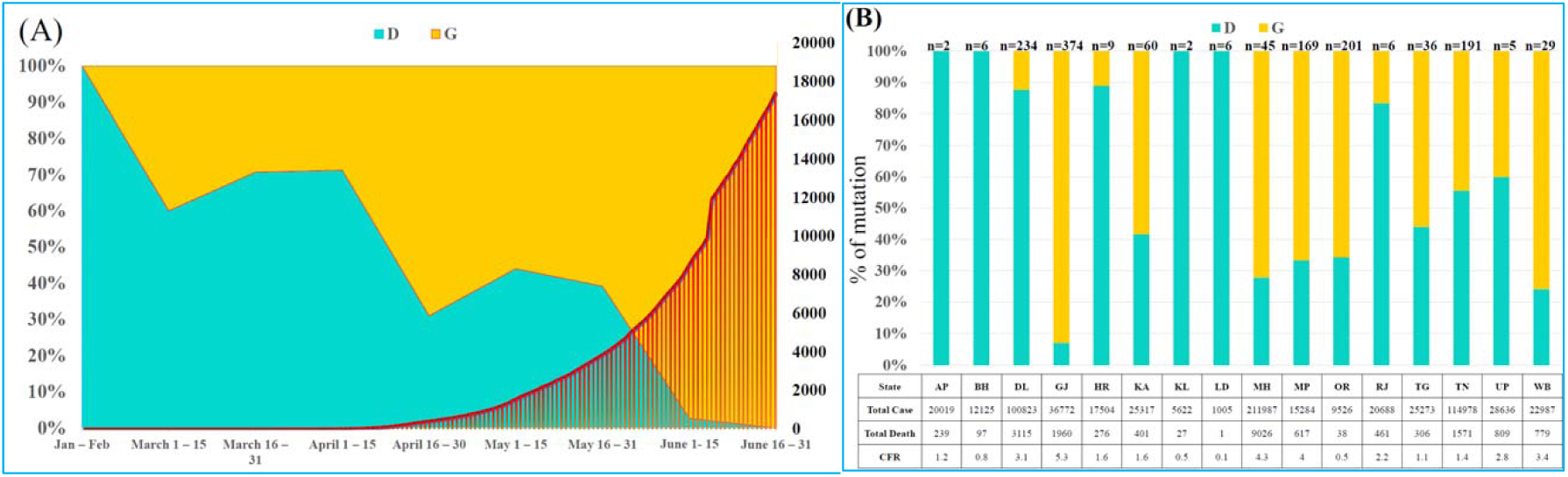
Temporal distribution of D614G genotype of SARS-CoV-2in India along with cumulative mortalities due to COVID-19 (Y1 axis represents percentage of D614G mutation and Y2 axis represent cumulative mortality). 3B. Distribution of D614G genotype in different state in India, total cases, mortality and case fatality rate in the states. [AP – Andhra Pradesh, BH – Bihar, DL – Delhi, GJ-Gujarat, HR – Haryana, KA – Karnataka, KL – Kerala, LD – Ladakh, MH – Maharashtra, MP – Madhya Pradesh, OR – Orissa, RJ – Rajasthan, TG – Telangana, TN – Tamil Nadu, UP – Uttar Pradesh, WB-West Bengal]

Phylogenetic tree based on amino acid sequences of spike protein for the selected isolates of India SARS-CoV-2were constructed using Nexstrain web tool (Fig 2). From the phylogeny it is apparent that, there are two major clusters of SARS-CoV-2in India based on the variation in the amino acid position 614. D614 is prominent in the reported cases of COVID-19 and newly detected G614 genotype circulating in the India similar to the situation in European countries (Pachetti et al., 2020). The transmission network analysis of the genomic data on Indian isolates of SARS-CoV-2shows that, the newly emerged G614 genotype that has reached India through travelers arriving from Europe and Middle East before establishing its prominent niche in the country (Fig 2). Overall observation made in this study suggests that G614 genotype has a pan India distribution and has steadily increased by replacing formerly introduced D614 genotype from Asian countries. But the distribution varied in different states in India (Fig 3). There were low numbers of sequences available from states like Andhra Pradesh (AP), Bihar (BH), Kerala (KL) and Ladakh (LD). Hence, we cannot draw any inference on the predominance of D614 in these states. But the difference between Delhi (DL) and Gujarat (GJ) is noteworthy. There were 214 sequences rom Delhi and 314 sequences from Gujarat. In Delhi, D614 predominated while in Gujarat, G614 predominated. Early hotspot in Delhi was a religious congregation involving travelers from Middle East as well as Asian countries and this could be reason for dominance of D614 in Delhi. But if S614 has a fitness advantage, it would have shifted the balance in favor of this genotype as pandemic advanced. This seems to be not the case at least in Delhi in India.

From the trends in data reported from India on COVID 19, it appears that CFR and incidence of G614 genotype is on the rise in the time path from the short to the long (Fig 3); but there is a paucity of evidences to prove the association among these factors. Korber et al (2020) noted that though viral loads were higher in infections with G614 genotype, there was no significant effect on hospitalization outcomes. Hence the study examines the probable association of D614G genotype with disease severity of COVID-19 across India through correlation analysis and the results are reported in Table 1. It appears that CFR is positively and strongly correlated with the increasing incidence of G614 genotype in different states of India. However, to predict the cause-effect relationship between CFR and level of G614 genotype, a liner regression model as presented in equation (1) has been used and the results are reported in Table 1.

**Table 1.** Model represents the association between D614G SNP in the SARS-COV2 genome and case fatality rate among the selected COVID-19 cases in India.

| A. Correlation Matrix |  |  |  |  |
| --- | --- | --- | --- | --- |
|  | CFR | D614G SNP |  |  |
| CFR | 1 | 0.619854 |  |  |
| D614G SNP | 0.619854 | 1 |  |  |
| B. OLS Regression Analysis |  |  |  |  |
| Dependent Variable: CFR |  |  |  |  |
| Particulars | Coefficient | Std. Error | t-Statistic | Prob. |
| C | 1.031447 | 0.482582 | 2.137353 | 0.0507 |
| Mutation | 0.027799 | 0.009406 | 2.955566 | 0.0104 |
| R-squared | 0.384219 | Mean dependent var |  | 2.118750 |
| Adjusted R-squared | 0.340235 | S.D. dependent var |  | 1.538059 |
| S.E. of regression | 1.249302 | Akaike info criterion |  | 3.399516 |
| Sum squared resid | 21.85059 | Schwarz criterion |  | 3.496090 |
| Log likelihood | -25.19613 | Hannan-Quinn criter. |  | 3.404462 |
| F-statistic | 8.735370 | Durbin-Watson stat |  | 2.054826 |
| Prob (F-statistic) | 0.010430 |  |  |  |

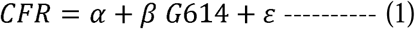

Where, CFR is COVID-19 case fatality ratio in India, α is the intercept of the equation, *β G*614 is beta coefficient of the deterministic variable of the model, ε is the error term. From the result, it appears that increasing incidence of G614 genotype reported from different states of India significantly contributes to higher fatality ratio among the COVID 19 positive cases. The beta of the regression coefficient is positive and statistically significant at 5% level. The model is a good fit as the r^2^ is at the acceptable level. It indicates that 38% changes in CFR are defined by the regressor variable in the model i.e. G614 genotype, whereas the remaining 62% changes in CFR are controlled by exogenous factors that are outside the present model. The F-statistic of the model is statistically significant at 1% level indicating that the results generated from the model are reliable. Hence the result from the regression model provides evidence to support the theoretical perception that CFR is adversely affected by G614 genotype thereby predicting that increasing trend in the G614 genotype may have profound negative impact on the clinical outcome of the COVID-19 cases in India.

## Conclusions

Even though there is a prima facie evidence on the infectivity potential of G614 mutant from the *in vitro* studies but how this mutation translates to clinical outcome of the COVID-19 needs to be addressed combining both laboratory experiments, clinical data and epidemiological investigations. In summary, this study unveils the genomic regions of genetic variation in the SARS-CoV-2 circulating in India, which is extensively dominated by G614 genotype with a strong correlation to CFR of COVID-19 posing enormous challenge for the effective prevention and management of COVID-19 cases in India.

## Acknowledgement

The authors are grateful to Nitte (Deemed to be University) for providing computing infrastructure for the execution of this research.

---

**Table S1.** List of mutation frequencies observed in genome of Indian SARS-CoV-2. The frequencies of mutations are calculated from 1383 SARS-CoV-2genomes.

| Sl. No | Protein mutation | SNP mutation | Type of mutation | Frequency | Name of the protein |
| --- | --- | --- | --- | --- | --- |
| 1 | NSP16:N298L | 21550 AA > CT | NonSynonymous mutation | 20 | 2'-O-ribose methyltransferase |
| 2 | NSP3:D455D | 4084 C > T | Synonymous mutation | 21 | Predicted phosphoesterase, papain-like proteinase |
| 3 | NSP12b:V871I | 16078 G > A | NonSynonymous mutation | 21 | RNA-dependent RNA polymerase |
| 4 | NSP3:P1103L | 6027 C > T | NonSynonymous mutation | 22 | Predicted phosphoesterase, papain-like proteinase |
| 5 | S:T1273T | 25381 A> C | Synonymous mutation | 23 | Spike |
| 6 | NSP4:A380V | 9693 C > T | NonSynonymous mutation | 27 | Transmembrane protein |
| 7 | NSP13:L130L | 16626 C> T | Synonymous mutation | 29 | Helicase |
| 8 | NSP4:D279N | 9389 G> A | NonSynonymous mutation | 36 | Transmembrane protein |
| 9 | NSP2:G339S | 1820 G > A | NonSynonymous mutation | 36 | Non-Structural protein 2 |
| 10 | NSP14:T372I | 19154 C > T | NonSynonymous mutation | 37 | 3'-to-5' exonuclease |
| 11 | NSP3:E545E | 4354 G > A | Synonymous mutation | 39 | Predicted phosphoesterase, papain-like proteinase |
| 12 | NSP3:S1285F | 6573 C > T | NonSynonymous mutation | 40 | Predicted phosphoesterase, papain-like proteinase |
| 13 | ORF3a:L46F | 25528 C > T | NonSynonymous mutation | 40 | ORF3a protein |
| 14 | NSP13:L92L | 16512 A > G | Synonymous mutation | 45 | Helicase |
| 15 | 3'UTR:29742 | 29742 G > A | Extragenic | 46 | NA |
| 16 | NSP3:S1197R | 6310 C > A | NonSynonymous mutation | 53 | Predicted phosphoesterase, papain-like proteinase |
| 17 | NSP3:V527V | 4300 G > T | Synonymous mutation | 56 | Predicted phosphoesterase, papain-like proteinase |
| 18 | NSP14:L177F | 18568 C > T | NonSynonymous mutation | 59 | 3'-to-5' exonuclease |
| 19 | S:T302T | 22468 G > T | Synonymous mutation | 60 | Spike |
| 20 | 3'UTR:29750 | 29750 C > T | Extragenic | 60 | NA |
| 21 | NSP2:Q496P | 2292 A > C | NonSynonymous mutation | 61 | Non-Structural protein 2 |
| 22 | NSP4:S76S | 8782 C > T | Synonymous mutation | 61 | Transmembrane protein |
| 23 | NSP4:F121F | 8917 C > T | Synonymous mutation | 62 | Transmembrane protein |
| 24 | N:S202N | 28878 G > A | NonSynonymous mutation | 66 | Nucleocapsid protein |
| 25 | NSP14:L495L | 19524 C > T | Synonymous mutation | 67 | 3'-to-5' exonuclease |
| 26 | ORF8:L84S | 28144 T > C | NonSynonymous mutation | 69 | ORF8 protein |
| 27 | S:L54F | 21724 G > T | NonSynonymous mutation | 73 | Spike |
| 28 | NSP3:N305N | 3634 C > T | Synonymous mutation | 78 | Predicted phosphoesterase, papain-like proteinase |
| 29 | NSP12b:N619N | 15324 C > T | Synonymous mutation | 78 | RNA-dependent RNA polymerase |
| 30 | NSP1:L16L | 313 C > T | Synonymous mutation | 146 | Leader protein |
| 31 | NSP3:A994D | 5700 C > A | NonSynonymous mutation | 147 | Predicted phosphoesterase, papain-like proteinase |
| 32 | NSP3:C39C | 2836 C > T | Synonymous mutation | 155 | Predicted phosphoesterase, papain-like proteinase |
| 33 | S:D294D | 22444 C > T | Synonymous mutation | 189 | Spike |
| 34 | N:S194L | 28854 C > T | NonSynonymous mutation | 191 | Nucleocapsid protein |
| 35 | N:RG203KR | 28881 GGG > AAC | NonSynonymous mutation | 269 | Nucleocapsid protein |
| 36 | M:Y71Y | 26735 C > T | Synonymous mutation | 275 | Membrane |
| 37 | NSP14:C279C | 18877 C > T | Synonymous mutation | 276 | 3'-to-5' exonuclease |
| 38 | ORF3a:Q57H | 25563 G > T | NonSynonymous mutation | 282 | ORF3a protein |
| 39 | NSP6:L37F | 11083 G > T | NonSynonymous mutation | 301 | Transmembrane protein |
| 40 | N:P13L | 28311 C > T | NonSynonymous mutation | 365 | Nucleocapsid protein |
| 41 | S:Y789Y | 23929 C > T | Synonymous mutation | 367 | Spike |
| 42 | NSP3:T1198K | 6312 C > A | NonSynonymous mutation | 383 | Predicted phosphoesterase, papain-like proteinase |
| 43 | NSP12b:A88V | 13730 C > T | NonSynonymous mutation | 405 | RNA-dependent RNA polymerase |
| 44 | NSP12b:P314L | 14408 C > T | NonSynonymous mutation | 800 | RNA-dependent RNA polymerase |
| 45 | Leader sequence<br>5'UTR:241 | C241T | Extragenic | 840 | NA |
| 46 | S:D614G | A23403G | NonSynonymous mutation | 845 | Spike |
| 47 | NSP3:F106F | 3037 C > T | Synonymous mutation | 853 | Predicted phosphoesterase, papain-like proteinase |

## References

1. Biswas NK, Majumder PP. Analysis of RNA sequences of 3636 SARS-CoV-2 collected from 55 countries reveals selective sweep of one virus type. Indian J. Med. Res. 2020 May 30.

2. Gao Y, Yan L, Huang Y, Liu F, Zhao Y, Cao L, Wang T, Sun Q, Ming Z, Zhang L, Ge J. Structure of the RNA-dependent RNA polymerase from COVID-19 virus. Science. 2020 May 15;368(6492):779–82.

3. Korber B, Fischer WM, Gnanakaran S, Yoon H, Theiler J, Abfalterer W, Hengartner N, Giorgi EE, Bhattacharya T, Foley B, Hastie KM. Tracking changes in SARS-CoV-2Spike: evidence that D614G increases infectivity of the COVID-19 virus. Cell. 2020 Jul 3.

4. Lin Q, Zhao S, Gao D, Lou Y, Yang S, Musa SS, Wang MH, Cai Y, Wang W, Yang L, He D. A conceptual model for the coronavirus disease 2019 (COVID-19) outbreak in Wuhan, China with individual reaction and governmental action. International journal of infectious diseases. 2020 Apr 1;93:211–6.

5. Liu Y, Gayle AA, Wilder-Smith A, Rocklöv J. The reproductive number of COVID-19 is higher compared to SARS coronavirus. Journal of travel medicine. 2020 Mar 1.

6. Mercatelli D, Triboli L, Fornasari E, Ray F, Giorgi FM. coronapp: a Web Application to Annotate and Monitor SARS-CoV-2Mutations. bioRxiv. 2020 Jan 1.

7. Mukherjee M, Goswami S. Global cataloguing of variations in untranslated regions of viral genome and prediction of key host RNA binding protein-microRNA interactions modulating genome stability in SARS-CoV2. bioRxiv. 2020 Jan 1. doi: https://doi.org/10.1101/2020.06.09.134585

8. Mukherjee M, Goswami S. Global cataloguing of variations in untranslated regions of viral genome and prediction of key host RNA binding protein-microRNA interactions modulating genome stability in SARS-CoV2. bioRxiv. 2020 Jan 1.

9. Pachetti M, Marini B, Benedetti F, Giudici F, Mauro E, Storici P, Masciovecchio C, Angeletti S, Ciccozzi M, Gallo RC, Zella D. Emerging SARS-CoV-2mutation hot spots include a novel RNA-dependent-RNA polymerase variant. Journal of Translational Medicine. 2020 Dec;18:1–9.

10. Potdar V, Choudhary ML, Bhardwaj S, Ghuge R, Sugunan AP, Gurav Y, Yadav PD, Shete A, Tomar S, Anukumar B, Kaushal H. Respiratory virus detection among the overseas returnees during the early phase of COVID-19 pandemic in India. Indian Journal of Medical Research. 2020 May 1;151(5):486.

11. Singh S, Sharma BB. Severe acute respiratory syndrome-coronavirus 2 and novel coronavirus disease 2019: An extraordinary pandemic. Lung India. 2020 May 1;37(3):268.

12. Somasundaram K, Mondal M, Lawarde A. Genomics of Indian SARS-CoV-2: Implications in genetic diversity, possible origin and spread of virus. medRxiv. 2020 Jan 1.

13. WHO, Coronavirus disease 2019 (COVID-19) situation report – 178, Data as received by WHO from national authorities by 10:00 CEST, 16 July 2020

14. Wu F, Zhao S, Yu B, Chen YM, Wang W, Song ZG, et al. A new coronavirus associated with human respiratory disease in China. Nature. 2020. https://doi.org/10.1038/s41586-020-2008-3

15. Zhang L, Jackson CB, Mou H, Ojha A, Rangarajan ES, Izard T, Farzan M, Choe H. The D614G mutation in the SARS-CoV-2 spike protein reduces S1 shedding and increases infectivity. bioRxiv. 2020 Jan 1.

